# Metabolic phenotyping of healthy and diseased human RPE cells

**DOI:** 10.1101/2024.02.28.582405

**Authors:** Saira Rizwan, Beverly Toothman, Bo Li, Abbi J. Engel, Rayne R. Lim, Sheldon Niernberger, Jinyu Lu, Cloe Ratliff, Yinxiao Xiang, Mark Eminhizer, Jennifer R. Chao, Jianhai Du

## Abstract

**Purpose:** Metabolic defects in the retinal pigment epithelium (RPE) underlie many retinal degenerative diseases. This study aims to identify the nutrient requirements of healthy and diseased human RPE cells.

**Methods:** We profiled nutrient utilization of various human RPE cells, including differentiated and dedifferentiated fetal RPE (fRPE), induced pluripotent stem cell derived-RPE (iPSC RPE), Sorsby fundus dystrophy (SFD) patient-derived iPSC RPE, CRISPR-corrected isogenic SFD (cSFD) iPSC RPE, and ARPE-19 cell lines using Biolog Phenotype MicroArray Assays.

**Results:** Differentiated fRPE cells and healthy iPSC RPE cells can utilize 51 and 48 nutrients respectively, including sugars, intermediates from glycolysis and tricarboxylic acid (TCA) cycle, fatty acids, ketone bodies, amino acids, and dipeptides. However, when fRPE cells lose their epithelial phenotype through dedifferentiation, nutrient utilization becomes restricted to 17 nutrients, primarily sugar and glutamine-related amino acids. SFD RPE cells can utilize 37 nutrients; however, compared to cSFD RPE and healthy iPSC RPE, they are unable to utilize lactate, some TCA cycle intermediates, and short-chain fatty acids. Nonetheless, they show increased utilization of branch-chain amino acids (BCAAs) and BCAA-containing dipeptides. Dedifferentiated ARPE-19 cells grown in traditional culture media cannot utilize lactate and ketone bodies. In contrast, nicotinamide supplementation promotes differentiation towards an epithelial phenotype, restoring the ability to use these nutrients.

**Conclusions:** Epithelial phenotype confers metabolic flexibility to healthy RPE for utilizing various nutrients. SFD RPE cells have reduced metabolic flexibility, relying on the oxidation of BCAAs. Our findings highlight the potentially important roles of nutrient availability and utilization in RPE differentiation and diseases.

## Introduction

Retinal pigment epithelium (RPE) is crucial in supporting photoreceptor function and survival through various functions, including the visual cycle, nutrient transport, light absorption, phagocytosis of outer segments, and formation of the blood-retina barrier^1^. To sustain these critical functions, the RPE relies on a robust mitochondrial metabolism by oxidizing fuels from the photoreceptors and choroidal supply^2^. Defects in RPE mitochondrial metabolism can cause epithelial-mesenchymal transition (EMT) and dedifferentiation, with loss of characteristic epithelial traits such as tight junctions, pigmentation and polarity, subsequently leading to photoreceptor death in retinal degenerative diseases, including Sorsby Fundus Dystrophy (SFD) and age-related macular degeneration (AMD)^2-4^.

Cultured human RPE cells can be valuable models to investigate RPE function and retinal disease, as they exhibit typical RPE morphology, function and signature gene expression, which closely resemble native RPE cells^5-8^. Human fetal RPE (fRPE), patient-derived induced pluripotent stem cells (iPSC) RPE, and immortalized ARPE-19 cells are commonly used RPE cultures. Human fRPE cells have been rigorously characterized in their physiology, gene expression and metabolism, considered the gold standard in RPE culture^5, 9, 10^. iPSC RPE have been well-characterized and have the advantage of modeling inherited retinal diseases from patients. SFD, a rare early onset macular degeneration, results from mutations in the tissue inhibitor of metalloproteinase-3 (TIMP3) gene^11^. In our previous study, we showed that iPSC RPE derived from SFD patients carrying the S204C mutation in *TIMP3* have irregular extracellular matrices (ECM) and basal laminar deposits, while correction of the S204C variant in SFD iPSC RPE (cSFD) attenuates these findings^12^. ARPE-19 cells offer advantages over fRPE and iPSC RPE in terms of easier availability and low cost. ARPE-19 cells show fibroblast-like morphology under traditional culture media. However, supplementation with nicotinamide (NAM), a precursor for NAD, can rapidly induce differentiation of ARPE-19 cells into cells with many epithelial characteristics through revitalization of mitochondrial metabolism^13, 14^.

In this study, we determined the metabolic phenotype of nutrient utilization in healthy and diseased human RPE. We conducted extensive metabolic screening of carbon and nitrogen sources across multiple human RPE cultures, including mature and dedifferentiated fRPE, healthy and SFD RPE, and ARPE-19 RPE cells under different culture media conditions. We found that healthy and mature RPE cells demonstrate robust metabolic flexibility, allowing them to utilize diverse nutrient sources. In contrast, dedifferentiated and SFD RPE cells have reduced metabolic flexibility, relying on specific nutrient sources. These cell-specific differences in nutrient utilization should provide insights into the underlying mechanisms of RPE differentiation and disease.

## Methods

### Reagent and key resources

All the reagents and key resources were detailed in Supplemental **Table S1** or methods.

### Human RPE cell culture

Human fRPE was isolated from human fetal eye cups with no identifiers, obtained from two distinct donors to the Birth Defects Research Laboratory at the University of Washington (UW) and cultured as previously described^9, 15^. RPE cells at passages 4-6 were used for experiments, and they were seeded at a density of either 5 × 10^4^ cells/well (regular density) or 5 × 10^3^ cells/well (low density) in 96-well plates in MEM RPE media (See details in **Table S1**). iPSC RPE, one SFD patient-derived iPSC RPE cells and CRISPR-corrected isogenic cSFD RPE cells were generated in a stepwise procedure as reported (see details in Supplemental Methods)^12^. Human ARPE-19 cells obtained from the American Type Culture Collection (ATCC) were used at passages 5-8 for the experiments. The cells were seeded at 2 × 10^4^ cells/well in 96-well plates and cultured in three different media to mimic dedifferentiated and differentiated states: 1) DMEM/F-12 media with 5% FBS, 2) MEM RPE media, and 3) MEM-NAM (MEM RPE media supplemented with 10 mM NAM as reported^13, 14^. All RPE cells were cultured for 4 weeks metabolic screening.

### Metabolic screening with Biolog Phenotyping MicroArrays

Biolog Phenotyping MicroArrays (Biolog Inc.) are 96-well plate assays that measure cellular metabolic activity in utilizing various carbon and nitrogen substrates. Each well contains a specific nutrient at 100 μM concentration, except for the negative control. Cells are added to the 96-well in Biolog’s proprietary nutrient-limiting media, followed by the addition of a proprietary redox dye. This dye is reduced by cellular NADH generated by substrate utilization, causing a color change measured by a plate reader. High absorbance indicates greater metabolic activity in utilizing specific substrate through NADH production. We used two Biolog Phenotype MicroArrays (PM-M1 for carbon sources and PM-M2 for nitrogen sources) to assess nutrient utilization in RPE cells (see details in **Table S1-3**). After pre-incubation with 60 uL of nutrient-limiting media (IFM 1, glutamine 200 μM, 1% dialyzed FBS, and 1% penicillin and streptomycin) for one hour, RPE cell-containing plates were washed with PBS and incubated with transferred pre-incubation media from PM-M1 or PM-M2 microarrays for 40 hours at 37^°^C in a 5% CO_2_ incubator. The redox dye mix MA (10 μL) was then added, and the plates were sealed and read at an absorbance of 595 nm every 15 minutes for 4 hours.

### ^13^C glutamine tracing in fRPE cells

Human fRPE cells seeded at regular and low densities were cultured for 4 weeks and changed into DMEM with 5 mM glucose and 1 mM ^13^C glutamine. After incubation for 48 hours, the RPE cells and media were collected for metabolite extraction and analysis of ^13^C glutamine-derived metabolites with gas chromatography mass spectrometry (GC MS) as previously reported^9, 10^.

### Statistics

All data are expressed as the mean ± standard deviation. Fold change of absorbance over negative control without nutrient source >1.5, or p<0.05 by Student’s unpaired two-tailed t-tests using GraphPad Prism 9.0, was considered significant. Heat maps were generated in Microsoft Excel.

## Results

### Mature human fRPE cells demonstrate remarkable metabolic flexibility in nutrient utilization

To study nutrient utilization in mature fRPE cells, fRPE were cultured for 4 weeks to maturity. RPE demonstrated characteristic cobblestone morphology and pigmentation (Supplemental **Fig S1A-C**). The mature RPE cultures were switched to nutrient-limiting media with or without a specific nutrient from the PM-M1 plate to screen the utilization of 91 carbon sources (**Fig 1A, Table S2**). Remarkably, mature RPE could utilize 23 carbon sources including sugars, glycolysis intermediates (such as sugar phosphates, lactate, and pyruvate), TCA cycle intermediates, nucleosides, fatty acids and ketone bodies (**Fig 1B-C**). In addition to glucose, mature RPE had the capacity to utilize other sugars such as fructose, mannose, and galactose (**Fig 1B-C**). Nucleosides, containing ribose moiety, can serve as alternative energy sources. Mature RPE could robustly utilize inosine, adenosine and uridine.

**Figure 1.**
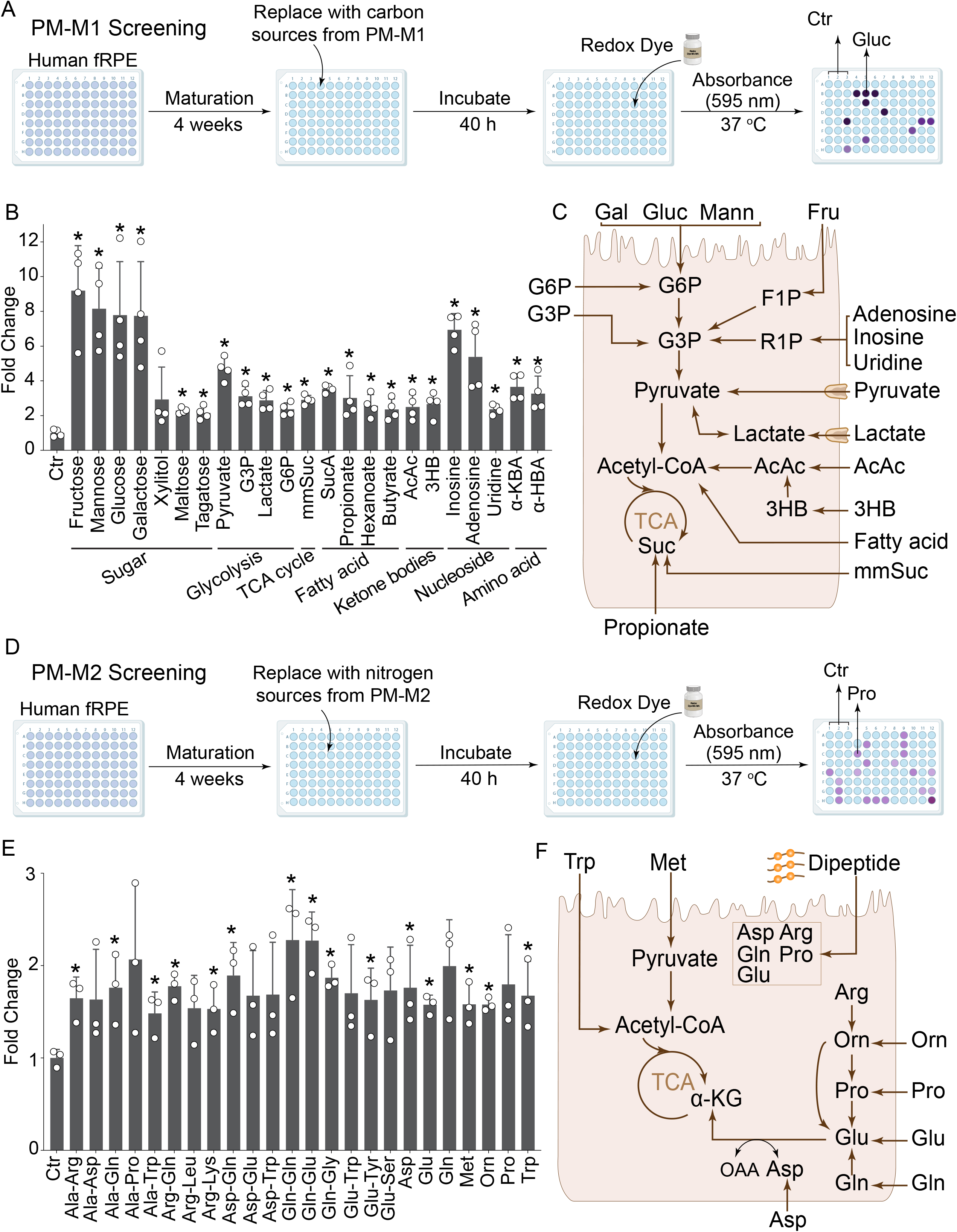
Metabolic phenotyping of mature human fRPE cells. (A) A schematic for metabolic phenotyping with carbon sources. Human fRPE cells were grown for 4 weeks in a 96-well plate and then switched into nutrient-limiting media containing different carbon sources in each well from PM-M1 plate. The utilization of nutrients leads to NADH production, casuing a color change in a redox sensitive dye to purple, which is quantified by a microplate reader at 595 nm. (B) The utilization of carbon sources by mature human fRPE cells and (C) an illustration of the metabolic pathways. (D) A schematic for metabolic phenotyping of nitrogen sources using PM-M2. (E) The utilization of nitrogen sources by mature human fRPE cells and (F) an illustration of the metabolic pathways. N=3. Fold change over negative control without nutrient source (Ctr)>1.5 or *P<0.05 over Ctr. Gluc, Glucose, Gal, galactose; Mann, Mannose; F1P, fructose 1-phosphophate; R1P, ribose 1-phosphate, G6P, glucose 3-phosphate; G3P, glycerol 3-phosphate; AcAc, Acetoacetic Acid; 3HB, β-Hydroxy Butyrate; MelA, Melibionic Acid.

Similar to PM-M1 carbon source screening, we evaluated the utilization of 92 nitrogen sources, including 27 amino acids and 60 dipeptides in mature RPE using the PM-M2 plate as illustrated in **Fig 1D**. Mature RPE could metabolize 7 amino acids and 17 dipeptides (**Fig 1E**). Except for methionine and tryptophan, the remaining 5 amino acids (proline, ornithine, glutamine, glutamate, and aspartate) follow a similar metabolic pathway in the generation of α-ketoglutarate (αKG) or oxaloacetate, which serve as fuels for the TCA cycle (**Fig 1E, 1F**). Notably, glutamine, glutamate, aspartate, and proline are also present in the 17 dipeptides. These findings suggest that mature fRPE cells have remarkable metabolic flexibility, enabling them to effectively utilize diverse nutrient sources to fuel their mitochondrial metabolism (**Fig 1C, 1E**).

### Dedifferentiated human fRPE cells lose metabolic flexibility, primarily relying on sugars and glutamine as their main nutrient sources

Seeding fRPE at low density induces their dedifferentiation into a fibroblast-like phenotype^5^. To understand nutrient utilization in dedifferentiated RPE cells, we seeded fRPE at 10% of regular density and cultured them for 4 weeks. As expected, these cells transitioned from their characteristic cobblestone structure to a fibroblast-like morphology, with upregulated expression of EMT markers including α-smooth muscle actin (αSMA) and vimentin (**Fig 2A**, Supplemental **Fig S1D-F, Fig S2**). Total protein content was similar in between mature and dedifferentiated RPE, indicating similar confluence between culture conditions (**Fig S3**). Compared to mature fRPE, dedifferentiated fRPE utilized fewer nutrients, with only 8 carbon and 6 nitrogen sources. Intriguingly, almost all carbon sources were sugar and sugar phosphates, while all nitrogen sources contained glutamine (**Fig 2A-C**). These results suggest that dedifferentiated RPE cells lose the ability to utilize multiple nutrient sources such as lactate, fatty acids, ketone bodies and proline. Instead, they become largely restricted to consuming sugars and glutamine (**Fig S4**). RPE seeded at low density for 1 week displayed a fibroblast-like morphology resembling that of 4-week cultured low-density fRPE cells but expressed less αSMA (**Fig S5A-B, Fig S1D-F, Fig S2**). Nutrient utilization in 1-week cells was largely consistent with those of 4-week fRPE, except for the capability to utilize galactose and adenosine at 1 week (**Fig S5B**), suggesting that time in culture has minimal impact on the metabolic phenotype of nutrient utilization in RPE cells seeded at low density. Consequently, we did not perform PM-M2 screening on 1-week fRPE.

**Figure 2.**
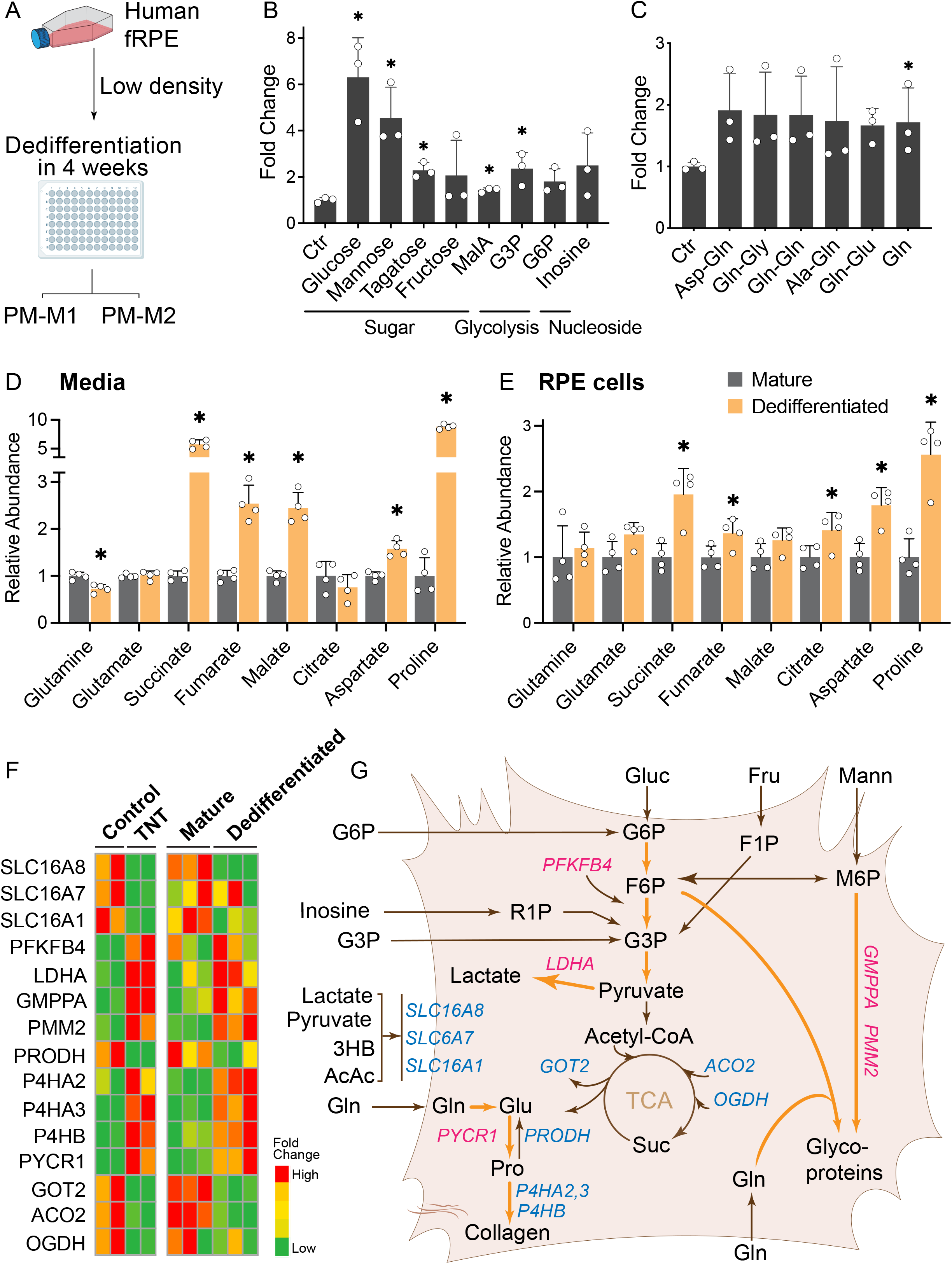
Metabolic phenotyping of dedifferentiated human fRPE. (A) Human fRPE were seeded at low density to induce dedifferentiation with a mesenchymal phenotype in a 96-well plate and then switched into nutrient-limiting media containing different carbon (PM-M1) or nitrogen sources (PM-M2) for metabolic phenotyping. (B) Nutrient utilization of carbon sources by dedifferentiated human fRPE. N=4. Fold change>1.5 or P<0.05 over Ctr. (C) Nutrient utilization of nitrogen sources by dedifferentiated human fRPE. N=3. Fold change>1.5 or P<0.05 over Ctr. (D-E) Dedifferentiated RPE consumes more glutamine for their mitochondrial metabolism. ^13^C-glutamine and its derived metabolites in media (D) and RPE cells (E) were analyzed by GC MS and presented as fold change of ^13^C labeled abundance over low density RPE. N=4. *P<0.05 vs mature RPE. (F) Heat maps of altered metabolic genes in dedifferentiated RPE cells induced by TNT (left panel) or multiple passage (right panel) (G) The dedifferentiated RPE cells lose the ability to utilize multiple nutrient sources but switch to use glutamine, enhancing proline and collagen synthesis. Magenta denotes an increase and blue denotes a decrease in gene expression or metabolic pathways. *PFKFB4*, 6-phosphofructo-2-kinase/fructose-2,6-bisphosphatase 4, *LDHA*, Lactate dehydrogenase A; *OGDH*, Oxoglutarate dehydrogenase; *PRODH*; Proline dehydrogenase; *PYCR1*, Pyrroline-5-carboxylate reductase 1; P4H4, Prolyl 4-hydroxylase subunit alpha or beta; *GMPPA*, GDP-mannose pyrophosphorylase A. *PMM2*, Phosphomannomutase 2.

To investigate how dedifferentiated RPE cells utilize glutamine, we incubated mature and dedifferentiated RPE with ^13^C glutamine for 48 hours and analyzed ^13^C glutamine-derived intracellular and extracellular metabolites (Fig 2D-E, Fig S6). Dedifferentiated RPE consumed more ^13^C glutamine in the media, which was utilized to produce mitochondrial intermediates and amino acids, especially succinate and proline (Fig 2D-E, Fig S6). These results suggest that dedifferentiated RPE cells have an impaired electron transport chain and reprogram their metabolism through reverse succinate dehydrogenase and proline dehydrogenase towards succinate and proline production.

We next analyzed the expression of metabolic genes related to nutrient utilization in two RNA-Seq data sets: 1) mature fRPE compared to dedifferentiated RPE induced by multiple passages^16^ and 2) mature primary adult RPE compared to dedifferentiated RPE induced by transforming growth factor-beta (TGFβ) and tumor necrosis factor α (TNFα)^17^. Both dedifferentiated RPE models had significant downregulation of genes involved in fatty acid oxidation, TCA cycle, transporters for lactate and ketone bodies, nucleotide metabolism, and amino acid metabolism, especially proline catabolism and glutamine synthetase (Fig 2F-G, **Table S4**). These data corroborate our findings that dedifferentiated RPE cannot efficiently utilize fatty acids, lactate, ketone bodies, proline and other nucleosides for mitochondrial metabolism. Importantly, genes involved in proline synthesis, glycolysis, collagen and heparin sulfate were substantially upregulated (**Fig 2F, Table S4**), suggesting dedifferentiated RPE cells utilize sugar and glutamine to activate proline synthesis and the hexosamine biosynthetic pathway to produce collagen and glycan for ECM remodeling **(Fig 2F)**.

### Human iPSC RPE cells have similar metabolic flexibility to fRPE cells but utilize more sugars and TCA cycle intermediates

To study nutrient utilization in human iPSC RPE cells, we cultured iPSC RPE to maturity at 4 weeks (**Fig S7**) and then replaced media with carbon or nitrogen sources from PM-M1 or PM-M2 for metabolic phenotyping (**Fig 3A**). iPSC RPE could utilize 31 carbon sources and 17 nitrogen sources (**Fig 3B-D**). Similar to fRPE, iPSC RPE demonstrated robust flexibility in utilizing various sugars, fatty acids, lactate, ketone bodies, nucleosides, amino acids, and dipeptides (**Fig 3B-D**). However, unlike fRPE, iPSC RPE showed a utilization preference for more sugars and TCA cycle intermediates and fewer amino acids and dipeptides (**Fig 3E-F**). These results suggest both healthy fRPE and iPSC RPE possess metabolic flexibility in adapting to different substrates, yet also have cell type-specific preferences for certain substrates.

**Figure 3.**
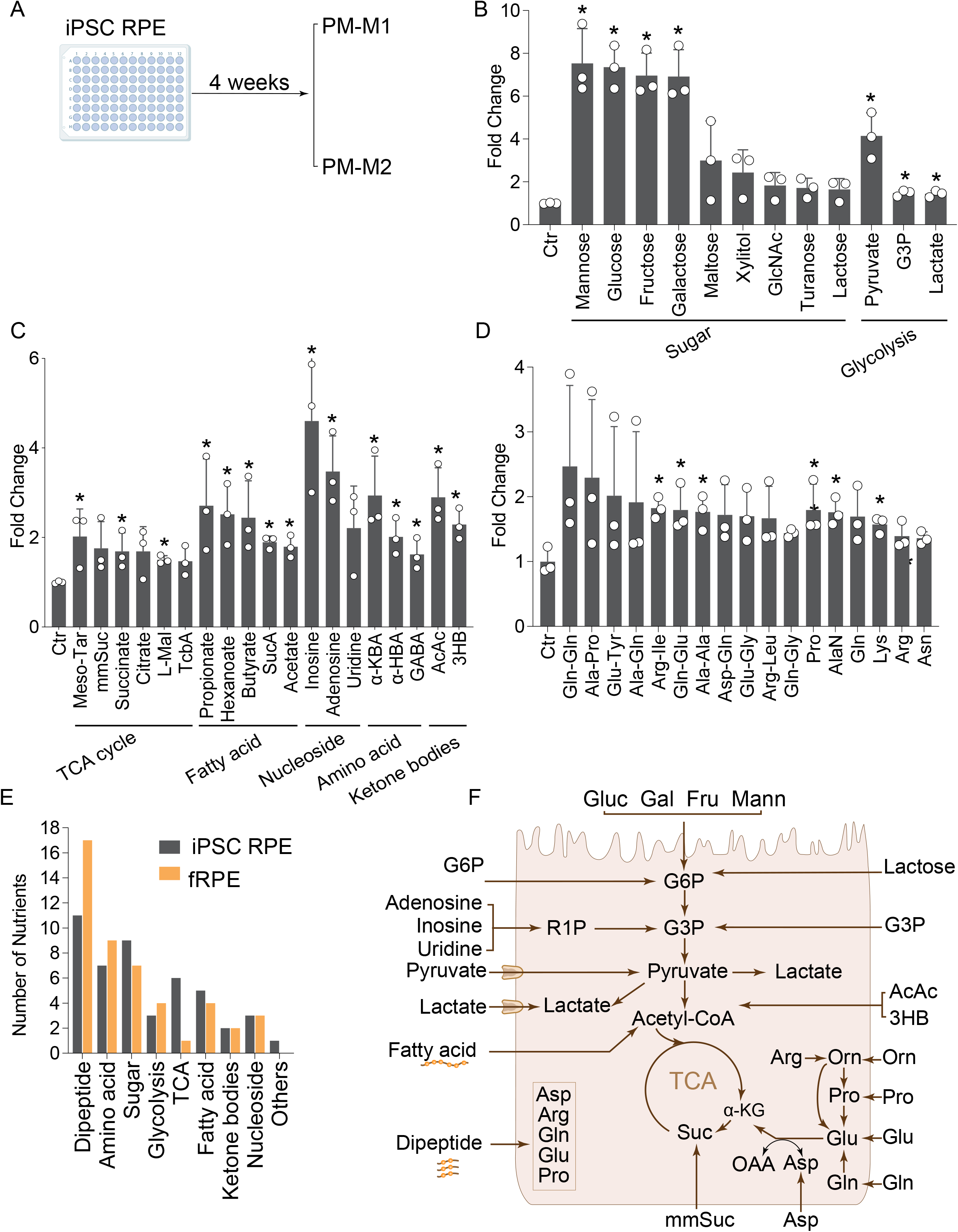
Metabolic phenotyping of healthy human iPSC RPE cells. **(A)** Human iPSC RPE cells were differentiated in RPE media for 4 weeks and switched into nutrient-limit media from PM1 or PM2 for metabolic phenotyping. (B-D) Nutrient utilization of carbon sources and nitrogen sources by human iPSC RPE cells. N=3. Fold change>1.5 or P<0.05 over Ctr. (E) A comparison of number of nutrients that were utilized between fRPE and iPSC RPE cells. (F) An illustration of nutrient utilization in iPSC RPE cells. GlcNAc, N-Acetyl-D-Glucosamine; Tcba, Tricarballylic Acid; AlaN, L-Alaninamide.

### SFD iPSC RPE cells utilize fewer intermediates from glycolytic and TCA cycle intermediates but more branch-chain amino acids (BCAAs)

We reported SFD iPSC RPE cells harboring the *TIMP3* S204 mutation have increased extracellular deposits and elevated intracellular 4-hydroxyproline^12^. These iPSC RPE and their isogenic controls (cSFD) were grown to maturity at 4 weeks before undergoing metabolic phenotyping with PM-M1 and PM-M2 (**Fig 4A**). Both SFD and cSFD had normal cobblestone structure and pigmentation (**Fig S8A-B**). SFD RPE utilized 19 carbon sources and 19 nitrogen sources, while cSFD RPE utilized 28 carbon sources and 7 nitrogen sources (**Fig 4B-E**). SFD RPE utilized different types of sugar, nucleosides, fatty acids, ketone bodies, amino acids and dipeptides (**Fig 4B, 4D**). However, SFD RPE could not utilize lactate, sugar phosphate, short-chain fatty acids (acetate and butyrate), succinate, αKG and gamma-aminobutyric acid (GABA). Notably, the CRISPR-corrected RPE cells restored the utilization of these nutrients (**Fig 4C-E**). SFD RPE could use proline and dipeptides containing glutamine, alanine and arginine. Intriguingly, many of the dipeptides utilized by SFD RPE cells contained BCAAs, leucine, isoleucine and valine, along with the ability to catabolize free leucine (**Fig 4D**). These metabolic phenotypes are different from fRPE and normal control iPSC RPE. While the cSFD RPE cells did not use the BCAAs, they also utilized significantly fewer nitrogen sources (**Fig 4E, Fig S8C**). These results suggest that SFD RPE cells have metabolic defects in recycling nutrients, such as lactate from the neural retina, and rely more on BCAAs, which may contribute to the formation of sub-RPE deposits of lipids and ECM proteins (**Fig 4F**).

**Figure 4.**
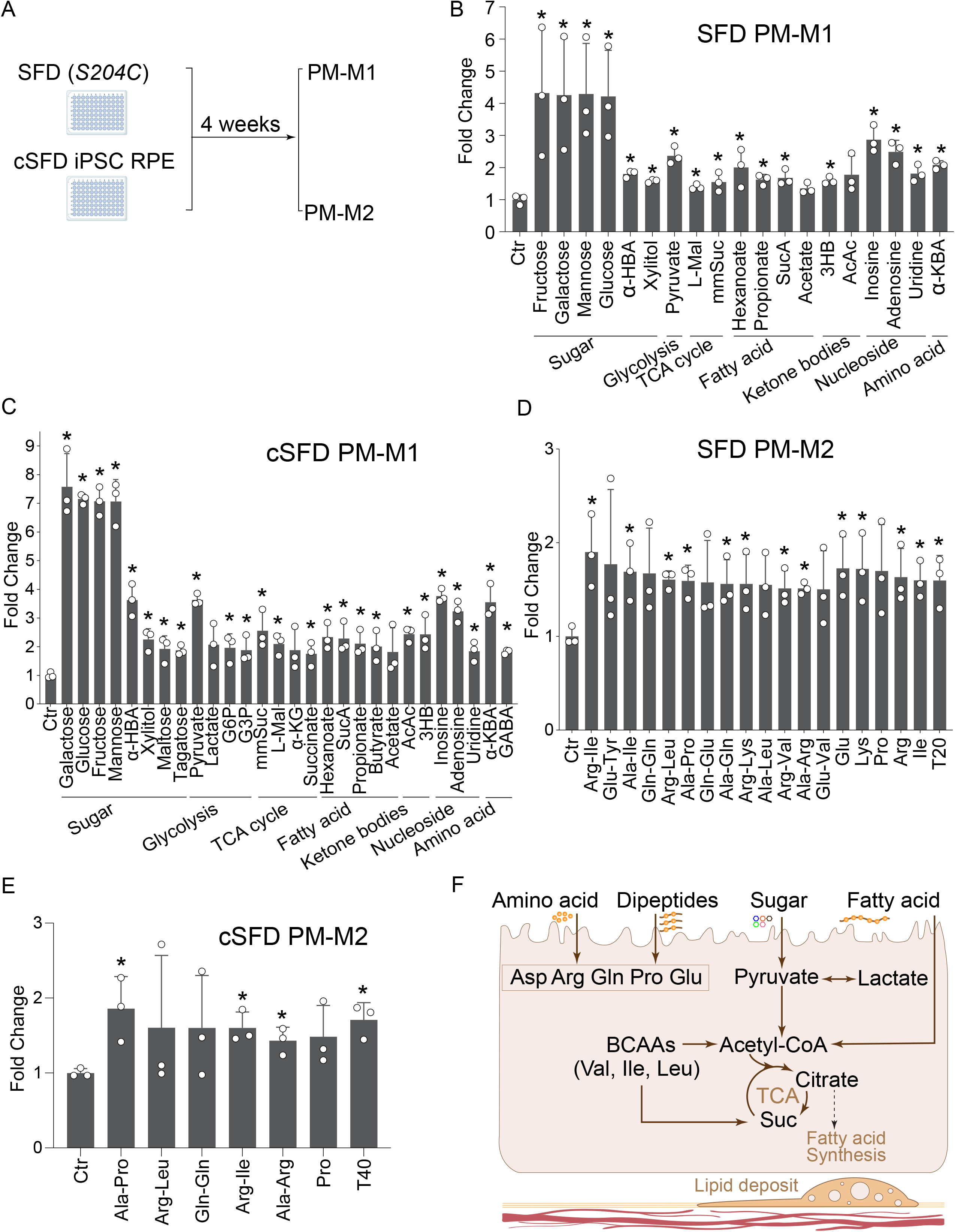
Metabolic phenotyping of IPSC SFD and cSFD RPE cells. (A) Human iPSC SFD RPE carrying TIMP3 S204C mutation and mutation corrected cSFD RPE cells were differentiated in RPE media for 4 weeks and then switched to nutrient-limit media with carbon or nitrogen sources from PM-M1 or PM-M2 for metabolic phenotyping. (B-E) Nutrient utilization of (B-C) carbon sources and (D-E) nitrogen sources in iPSC SFD RPE cells. N=3. Fold change>1.5 or P<0.05 over Ctr. (F) An illustration of altered nutrient utilization in SFD iPSC RPE cells. L-Mal, L-Malic Acid; mmSuc, mono methyl succinate; 3HB, β-Hydroxy Butyrate; BCAAs, Branch-Chained Amino Acids.

### Culture media composition controls the differentiation and nutrient utilization in ARPE-19 cells

ARPE-19 cells, typically cultured in DMEM/F-12 media per manufacturer’s instructions, undergo rapid differentiation into a mature RPE-like phenotype grown MEM-α-based RPE culture media supplemented with NAM, ^13, 14^. To investigate the influence of media composition on nutrient utilization, we cultured ARPE-19 cells under three media (DMEM/F12, MEM RPE media and MEM RPE media plus NAM) for 4 weeks and focused only on the screening of carbon sources using the PM-M1 plate (**Fig 5A**). DMEM/F-12 cultured cells (fibroblast-like morphology, **Fig S9A**) utilized 16 carbon sources (**Fig 5B**) but, similar to dedifferentiated fRPE, could not metabolize lactate, ketone bodies, or xylitol (**Fig 5B, Fig 6A**). MEM RPE media partially differentiated ARPE-19 cells (cobblestone structure, **Fig S9B**) and increased their carbon source utilization to 24 (covering 94% of those used by DMEM/F12 cells, Fig 5C). While they gained the ability to utilize ketone bodies and some sugars, lactate and xylitol remained unutilized (**Fig 5C**). Supplementation with NAM in MEM RPE media induced mature RPE-like morphology (**Fig S9C**) and allowed ARPE-19 cells to utilize lactate, ketone bodies, sugar phosphates, xylose, and tagatose (**Fig 5D-E**). This metabolic phenotype resembles that of mature fRPE and iPSC RPE, although ARPE-19 cells reduced utilization of short-chain fatty acids and maltose (**Fig 6A**). These findings highlight the critical role of nutrient availability and utilization in RPE differentiation.

**Figure 5.**
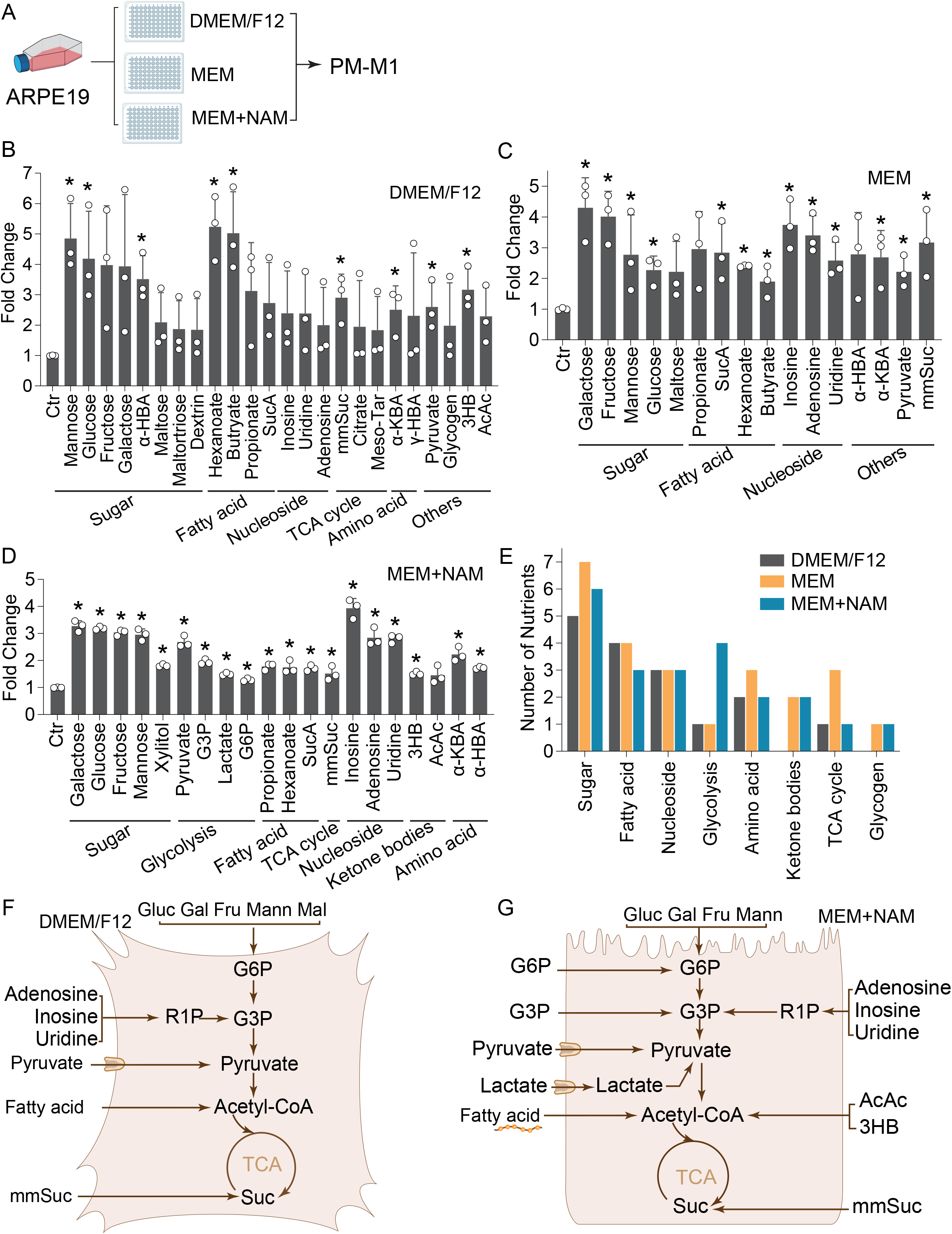
Metabolic phenotyping of ARPE-19 cells cultured in different media. (A) ARPE-19 cells were cultured in three different media: DMEM/F-12, MEM RPE media, and MEM RPE media with 10 mM NAM for 4 weeks. All the cells were then changed into nutrient-limit media with carbon sources from PM-M1 for metabolic phenotyping. (B-D) Nutrient utilization of carbon sources of ARPE-19 cells cultured in (B) DMEM/F12 (C) MEM RPE media, (D) MEM RPE media with NAM. Fold change>1.5 or P<0.05 over Ctr. (E) A comparison of number of nutrients that were utilized in ARPE-19 cells grown in different culture media. (F-G) An illustration of altered nutrient utilization in APRE-19 cells cultured between DMEM/F12 and MEM RPE media with NAM. Meso-Tar, Meso Tartatic acid; mmSuc, mono methyl succinate; AcAc, Acetoacetic Acid; 3HB, β-Hydroxy Butyrate.

**Figure 6.**
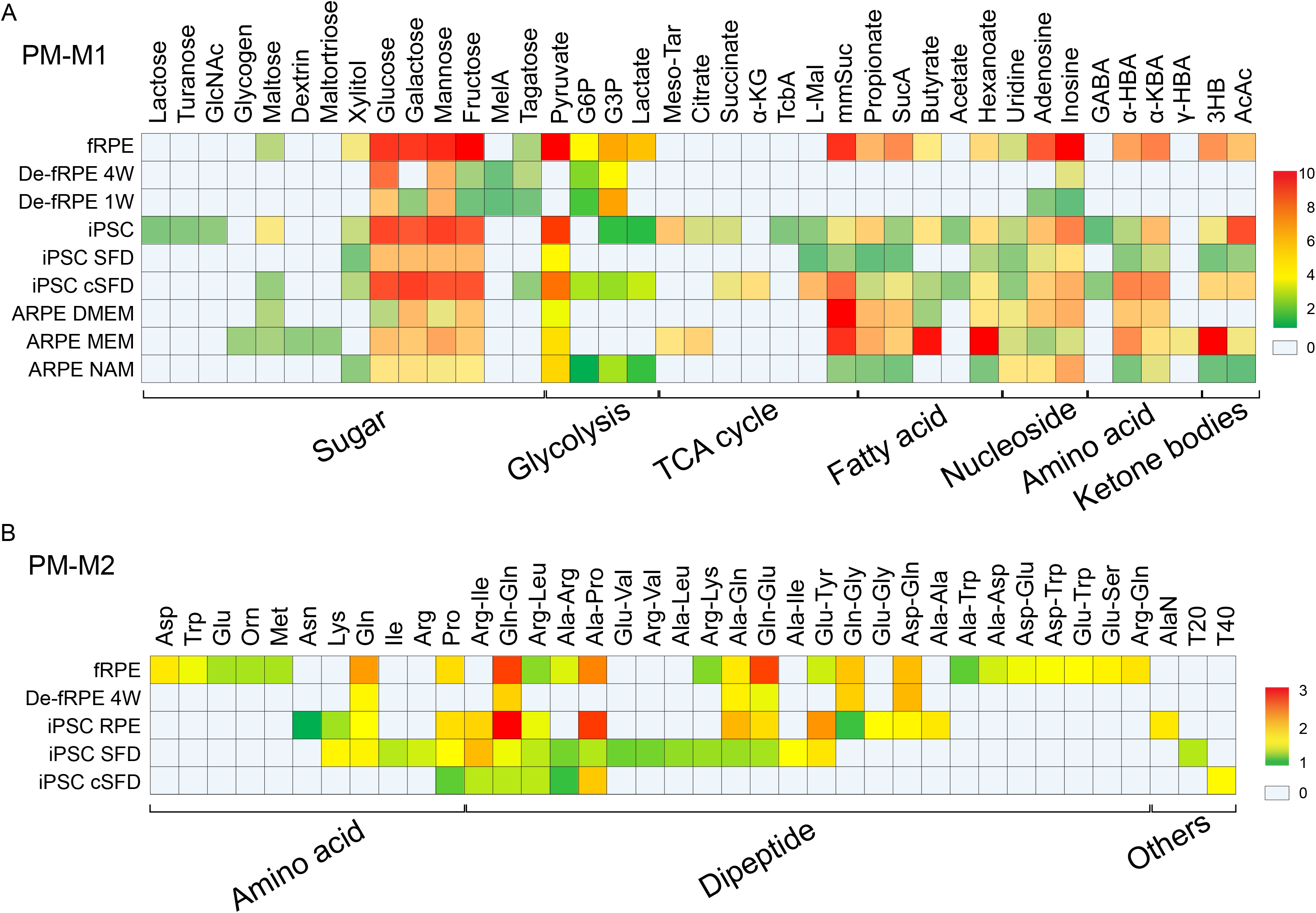
Comparison of metabolic phenotyping profiles of human fRPE cells, dedifferentiated fRPE, iPSC RPE cells, SFD RPE, cSFD RPE and ARPE-19 cells grown under different culture media. Heat map visualization of nutrients that were differentially utilized by different RPE cells from (A) PM-M1 nutrients and (B) PM-M2 nutrient microarray plates. De-fRPE, dedifferentied fRPW with low density seedings; ARPE, ARPE-19 cells; MelA, Melibionic Acid; GlcNAc, N-Acetyl-D-Glucosamine; Tcba, Tricarballylic acid; mmSuc, mono methyl succinate; AlaN, Alaninamide.

## Discussion

In this study, we identified distinctive metabolic phenotypes of nutrient utilization in healthy and diseased human RPE cells (**Fig 6**). Mature fRPE and iPSC RPE are metabolically flexible, capable of adapting to various nutrient sources. Conversely, dedifferentiated RPE cells and SFD patient-derived iPSC RPE cells have reduced flexibility and are biased towards fewer, specific substrates. Our findings suggest the critical roles of nutrient utilization in RPE differentiation and dedifferentiation.

The RPE must manage a substantial nutrient influx originating daily from phagocytosed lipid- and protein-rich photoreceptor outer segments, “waste products” such as lactate produced by the neural retina, and nutrient-laden blood through the choroidal circulation^1, 2^. In addition to the direct transport of nutrients to the neural retina, RPE can recycle and generate nutrients via lysosomes and mitochondria to support retinal metabolism, thereby establishing a metabolic ecosystem between the outer retina and RPE^1, 2, 18^. The ability to utilize a variety of nutrients is crucial for maintaining this metabolic ecosystem, and disruption of this balance due to mitochondrial and/or lysosomal dysfunction in RPE can lead to retinal degeneration^3, 4, 19^. Healthy human RPE cells are known to be versatile in utilizing different nutrients, including glucose, galactose, lactate, fatty acids, succinate, proline, glutamine and other amino acids^9, 18, 20-23^. Our results confirm this versatility, demonstrating that both healthy fRPE and iPSC RPE cells are capable of utilizing these nutrients.

Furthermore, we found that human RPE cells utilize a wide range of other nutrients including various sugars, ketone bodies, nucleosides and dipeptides. Mannose and galactose, for example, are components of glycoproteins found in abundance in outer segment proteins such as rhodopsin and peripherin 2^24-26^. The ability to utilize these sugars from the daily degradation of outer segments may allow RPE to preserve more glucose to supply for photoreceptors. Additionally, like the liver, human RPE cells can synthesize ketone bodies, which can be utilized by the neural retina^21, 27^. Interestingly, we found that mature human RPE, but not dedifferentiated RPE cells, can readily utilize ketone bodies, suggesting that the ability to utilize ketone bodies might be a characteristic of RPE metabolism. Consistent with this metabolic capability, mature RPE expresses key genes involved in ketone body degradation^21^, a feature distinct from the liver, which cannot utilize ketone bodies. Our findings underscore the metabolic flexibility of healthy mature RPE in nutrient utilization, which is an essential function for RPE to support retinal metabolism and health.

RPE EMT and dedifferentiation have been implicated in the pathogenesis of proliferative vitreoretinopathy and AMD^28^. We found dedifferentiated RPE cells were unable to utilize alternative fuels like lactate, fatty acids, ketone bodies and proline, but, instead, prefer to utilize sugars and glutamine. RNA-seq data from EMT models show reduced expression of genes involved in the TCA cycle, lactate transport, ketone body utilization and proline catabolism and glutamine synthesis. In contrast, gene expression associated with glycolysis, proline synthesis, collagen synthesis and glycoproteins are upregulated (**Fig 2E, Table S4**). Consistently, dedifferentiated RPE cells have been reported to switch from oxidative phosphorylation to glycolysis and lose the ability to utilize proline as a fuel^5, 15^. Inhibition of mitochondrial metabolism in human RPE cells or mouse RPE is sufficient to cause the activation of glycolysis, RPE dedifferentiation and retinal degeneration^4, 29^. Furthermore, mitochondrial respiration is diminished in cultured primary RPE cells from AMD donors^30^. Inhibiting mitochondrial respiration impedes proline consumption while increasing glucose consumption and lactate production^31^. The inability of affected RPE to utilize exogenous lactate and other alternative nutrient sources may be an important mechanism in AMD pathogenesis.

A key feature in dedifferentiated cells is their overproduction of ECM proteins, primarily collagens and glycoproteins, through activation of cytokine signaling such as TGFβ and TNFα^32, 33^. In fibroblasts, TGFβ stimulates the oxidation of glucose and glutamine to synthesize proline and other amino acids crucial for ECM protein production^34^. Interestingly, proline synthesis acts as a vent for growth when cells are under mitochondrial redox stress or hypoxia by recycling NADH into NAD^+^, thus enhancing the oxidation of glutamine and glucose^34, 35^. Proline synthesis may be particularly crucial in dedifferentiated RPE cells, given their inhibited mitochondrial metabolism and heavy reliance on lactate production for NAD^+^ regeneration. Similarly, to compensate for the deficiency in the electron transport chain or hypoxia, fumarate can serve as an electron acceptor to produce succinate^36^, which is accumulated in dedifferentiated RPE. Strategies aimed at relieving mitochondrial redox stress and reverting proline synthesis to proline catabolism or succinate production could hold promise for treating RPE dedifferentiation.

*TIMP3*, a risk gene for AMD, is secreted by the RPE and plays a critical role in ECM remodeling by inhibiting matrix metalloproteinases^37^. Specific mutations in *TIMP3* result in SFD, which is characterized by a thickened layer of basal laminar drusen-like sub-RPE deposits containing lipids and ECM proteins^12,38^. Unlike healthy or mutation-corrected RPE cells, SFD RPE cells utilize more free BCAAs or dipeptides containing BCAAs (**Fig 4F, Fig 6**). Leucine and isoleucine are ketogenic amino acids, and their oxidation promotes the synthesis and transport of fatty acids and cholesterol^39, 40^. BCAA oxidation is closely associated with dysregulated glucose and lipid metabolism in diabetes and non-alcoholic fatty liver diseases^39, 41, 42^. Interestingly, intermediates of the BCAA degradation pathway are significantly elevated in the plasma of AMD patients^43^, and *BCAT1*, a key gene in BCAA degradation, is upregulated in the RPE/choroid from AMD donors^44^. Moreover, in addition to proline, ECM proteins are highly enriched in BCAAs^45^. Our findings suggest that SFD RPE may upregulate the BCAA degradation pathway by utilizing BCAAs from free amino acids or protein degradation, contributing to the formation of sub-RPE deposits.

There are a few limitations in this study. We did not normalize nutrient utilization to specific cell numbers, although total cellular protein content was similar between dedifferentiated and mature RPE by 4 weeks in culture. Nutrient utilization in additional dedifferentiation models need to be conducted in future studies. Separately, high absorbance reflects greater reduction of NADH generated from a specific nutrient. The Biolog assays cannot detect the reduction of FADH2 or other metabolic pathways such as the synthesis of amino acids, nucleotides and fatty acids. In addition, the metabolic phenotyping assay can only measure the capability and capacity of utilizing a specific nutrient but not preferences of nutrient utilization because only one nutrient is available at a time. Therefore, the utilization might be different *in vivo*, and additional functional studies with nutrient supplements or genetic manipulations of metabolic pathways are needed to interpret these findings. Finally, testing additional iPSC RPE lines from a larger cohort of normal and affected donors will help control for variability in genetic and environmental backgrounds.

In conclusion, metabolic phenotyping offers a sensitive and powerful platform for measuring nutrient utilization in healthy and diseased human RPE cells. The distinct patterns of nutrient utilization of various RPE cells provide valuable insights into the mechanisms of RPE dedifferentiation and disease.

## Acknowledgments

This work was supported by National Institutes of Health Grant EY034364 (J. R. C. and J. D), EY03459 (J. R. C. and J. D), EY031324 (JD), EY032462(JD), the Retina Research Foundation (JD), and funds for Core facilities P20 GM103434 and P20 GM144230 (WV INBRE grant). Fetal RPE tissue samples were made available through Dr. Ian Glass and the Birth Defects Research Laboratory at the University of Washington (UWR24HD000836).

