## Supplementary Information for "Metabolic phenotyping of healthy and diseased human RPE cells"

This file includes:

Supplementary Methods

Supplementary Tables S1-3

Supplementary Figures S1-9

### Supplementary Methods

#### *Human fetal RPE culture*

RPE was isolated from human fetal eye cups with no identifiers, obtained from donors to the Birth Defects Research Laboratory at the University of Washington (UW) and cultured as previously described[1,2]. All procedures involving human tissue were done under an approved protocol by Institutional Review Boards of the University of Washington and West Virginia University and conformed to the ethical principles outlined in the Declaration of Helsinki. Two fetal RPE lines from distinct donors were used. RPE cells at passages 4-6 were used for experiments, and they were seeded at density of either 5 x $10^4$ cells/well (regular density) or 5 x $10^3$ cells/well (low density) in 96-well plates. Cells were grown for 4 weeks in MEM RPE media (See details in Table S1) at regular density or 1-4 weeks at low density to induce dedifferentiation before metabolic screening.

#### *Generation and culture of normal human iPSC RPE, SFD iPSC RPE, and cSFD iPSC RPE*

Informed consent was obtained from all subjects prior to inclusion in the study (University of Washington IRB-approved STUDY00010851). Peripheral blood mononuclear cells were isolated from one control donor without retinal degeneration and one SFD patient to generate patient-derived iPSC RPE cells in a stepwise procedure as reported (see details in Supplemental Methods)[3]. CRISPR-edited cSFD RPE were generated by eletroporating SFD RPE cells with Cas9-gRNA (0.75µM, GGGGCCTATTTTCGTAGTAG, Synthego) ribonucleoprotein (RNP) complex and ssDNA donor (4µM, IDT) using Amaxa Nucleofector (Human Stem Cell Kit 2). DNA was isolated from individual iPSC colonies and sequenced to identify the gene-edited iPSC colony. To assess potential off-target effects resulting from CRISPR-editing, off-target sites were identified using CRISPOR (http://crispor.tefor.net), which revealed two intronic sites with three mismatches. These regions were sequenced in the gene-edited iPSC RPE, and no genetic alterations were detected[3]. All iPSC and gene-edited iPSC RPE cells were maintained in MEM RPE media, detached with 0.25% Trypsin-EDTA and seeded at 5x$10^4$ cells/well into 96-well plates. RPE cells at passages 4-6 were cultured for 4 weeks before metabolic screening.

#### *ARPE-19 cell culture*

Human ARPE-19 cells obtained from the American Type Culture Collection (ATCC) were used at passages 5-8 for the experiments. The cells were seeded at 2 x 104 cells/well in 96-well plates and cultured in three different media to mimic dedifferentiated and differentiated states: 1) DMEM/F12 media with 5% FBS, 2) MEM RPE media, and 3) MEM-NAM (MEM RPE media supplemented with 10mM NAM as reported[4,5]. All the ARPE-19 cells were cultured for 4 weeks before metabolic screening.

#### *Immunofluorescence*

RPE seeded in Matrigel-coated 8well chambers at low or high densities were fixed after 1 or 4 weeks with 4% PFA, and permeabilized with 0.1% triton X-100. Blocking was done using 5% bovine serum albumin for 1 hour at room temperature, and primary antibodies (**Table S1**) were incubated overnight at 4$^{o}$C. Following three PBS washes, secondary antibodies were incubated for 1 hour at room temperature, followed by fluorophore conjugated phalloidin for 30 min. Cells were counterstained with DAPI and mounted for confocal microscopy. Images were taken using a 20X objective on the Leica DM6000 CS confocal microscope (Leica Microsystems).

***Protein concentration assays***

Fetal RPE were seeded on Matrigel-coated 96well plates at a total cell count of either 5 x $10^5$ or 0.5 x $10^5$ cells per well. To quantify protein content, 4-week-old RPE were scraped in Mammalian Protein Extraction Reagent (MPER) with Halt protease inhibitor (ThermoFisher). n=4 of a 96well plate were pooled into a single tube. Protein lysate was sonicated, spun down to remove cell debris, and quantified using Bradford protein assay (Pierce, ThermoFisher) against a protein standard curve. Data shown as mean ± SEM.

***Analysis of gene expression data in healthy and dedifferentiated RPE cells***

We analysed two RNA-Seq data sets: 1) mature fRPE vs dedifferentiated RPE induced by multiple passages[6] and 2) mature primary adult RPE vs dedifferentiated RPE induced by transforming growth factor-beta (TGFβ) and tumor necrosis factor α (TNFα)[7]. In the first data set, passage 0 (P0) and passage 5 (P5) fRPE were cultured for 32 days before RNA-Seq. The P0 RPE cells had typical epithelial structure, but P5 was dedifferentiated into fibroblast-like morphology. We compared the expression of metabolic genes in P5 compared to P0 and their fold change was calculated. For the second data set, primarily cultured adult RPE cells were stimulated with TGFβ + TNFα (TNT) to induce dedifferentiation. The metabolic gene expression of TNT vs control RPE was presented as fold change. All the gene expression data are in FPKM (fragments per kilo base per million mapped reads). The fold change was colored based on the level of expression.

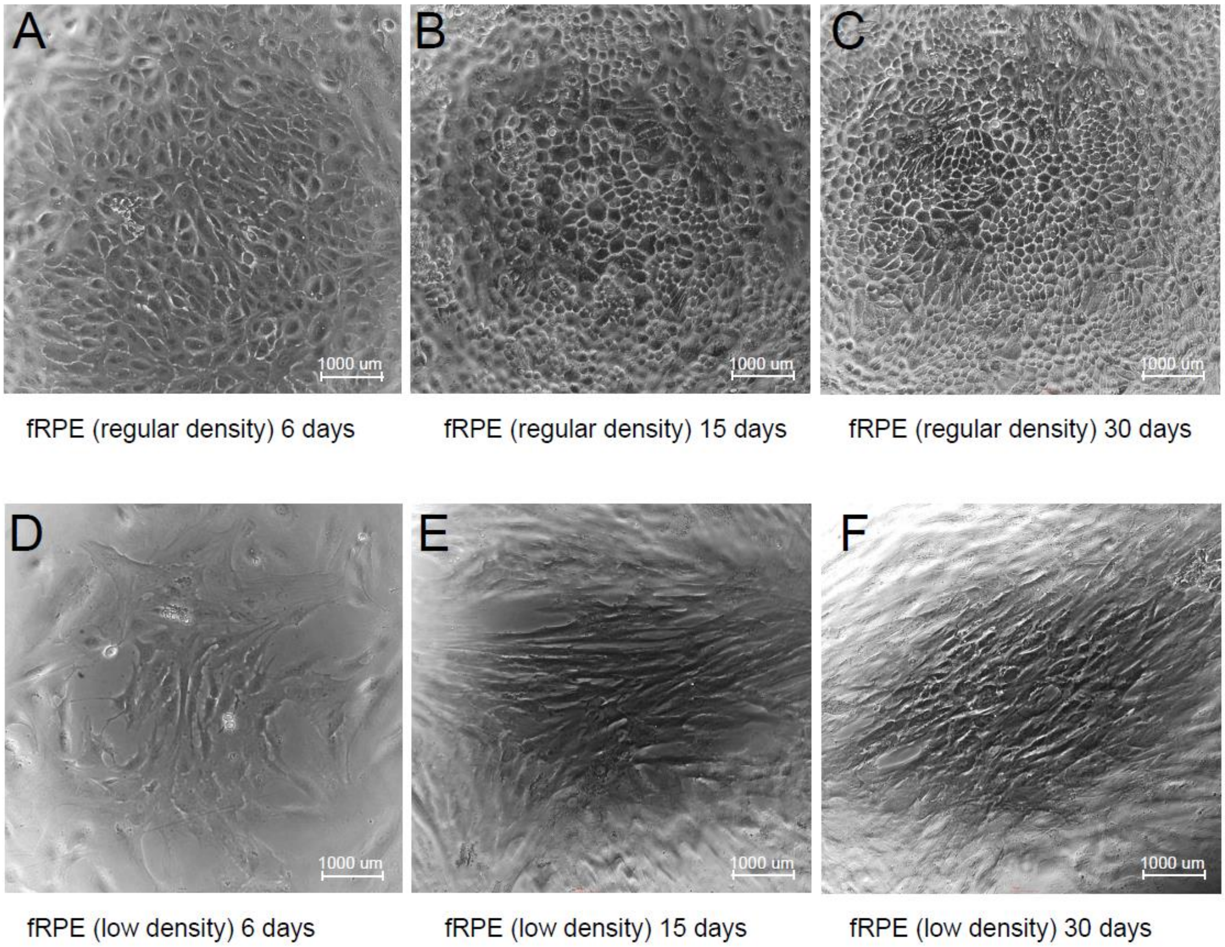


**Supplementary Figure S1. Morphology of human fetal RPE cells seeded at regular and low density**. (A-C) Representative bright-field images of human fetal RPE cells grown in a 96-well plate after seeded at regular density 5 x $10^5$ cells per well for different days. (D-E) Representative images of human fRPE cells seeded at low density 0. 5 x $10^5$ cells per well in a 96-well plate. Scale = 1000µm.

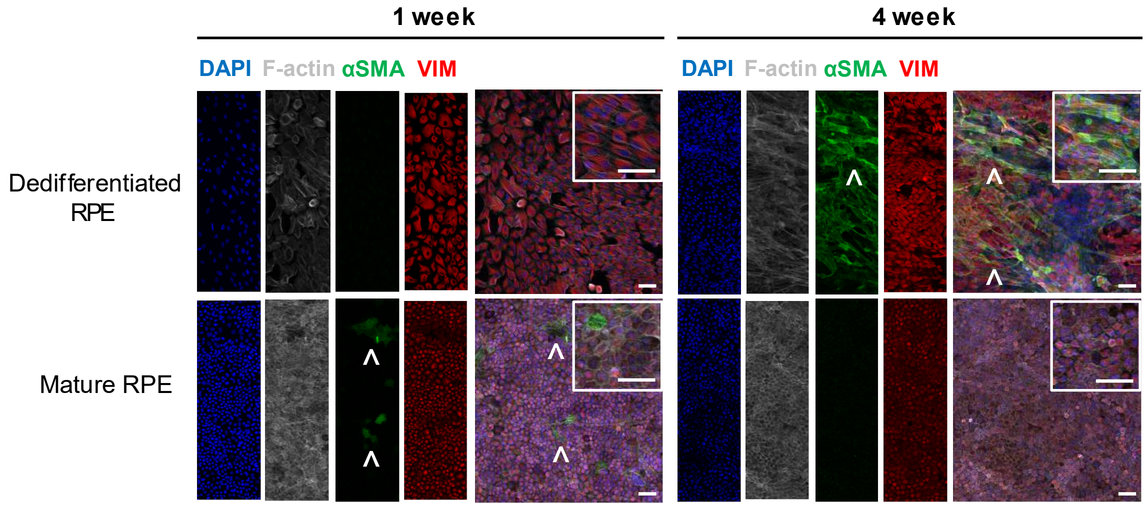


**Supplementary Figure S2. Immature and dedifferentiated RPE express αSMA.** RPE seeded in Matrigel-coated 8well chambers at low or regular density identical to that of 96well plates were fixed with 4% PFA and stained after 1 or 4 weeks in culture. Representative immunofluorescent images are shown with F-actin (Invitrogen A12380) in grey, αSMA (Abcam ab7817) in green and Vimentin (Cell Signaling Tech 5741) in red. αSMA positive cells (^) were seen in 1-week immature RPE seeded at regular density but were surprisingly absent in dedifferentiated RPE. By week 4, mature RPE had little to none αSMA positive cells, while the majority of dedifferentiated RPE cells were stained with αSMA. Vimentin was seen in all cells at all maturities, however staining intensity was much higher in low density cells, as well as αSMA positive cells that underwent EMT. Scale = 50µm.

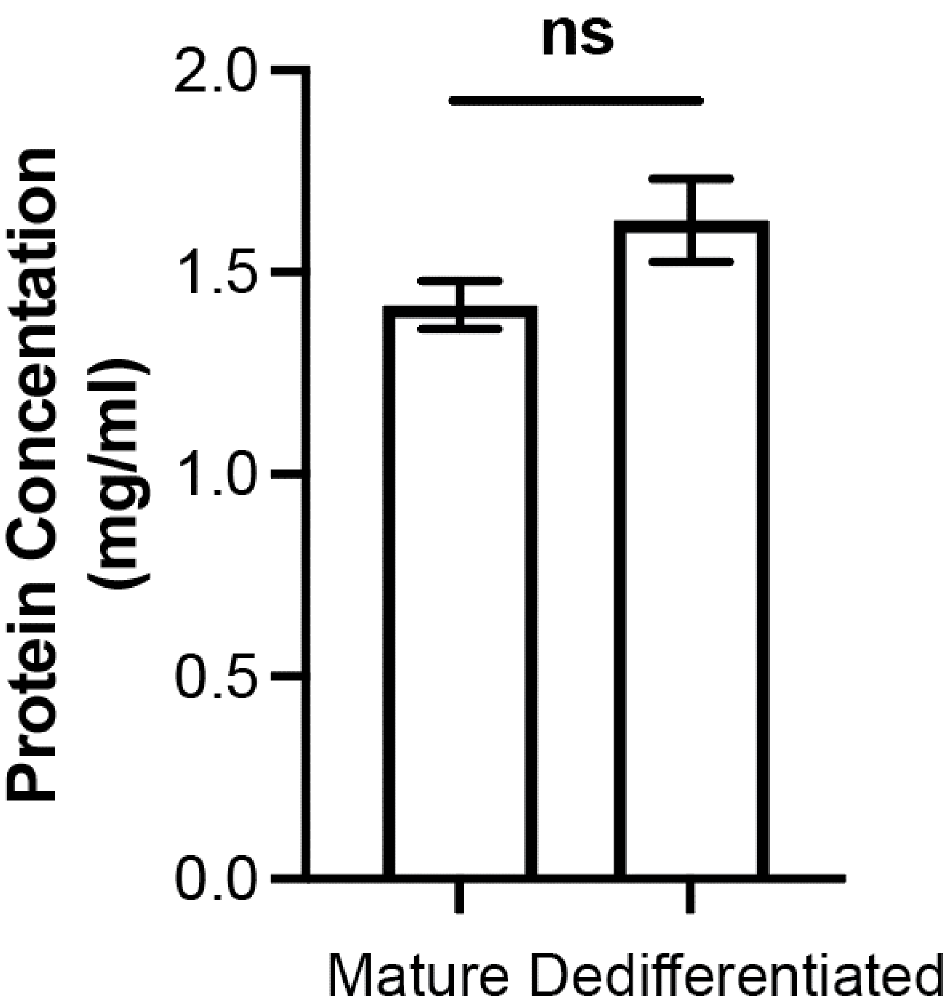


**Supplementary Figure S3. Dedifferentiated RPE cells have similar protein content to the mature RPE cells.** Fetal RPE were seeded on Matrigel-coated 96well plates at a total cell count of either 5 x $10^5$ or 0.5 x $10^5$ cells per well for mature and dedifferentiated RPE respectively. After culture for 4 weeks, RPE cells were scraped, lysed, and quantified for protein concentrations. Data shown as mean ± SEM.

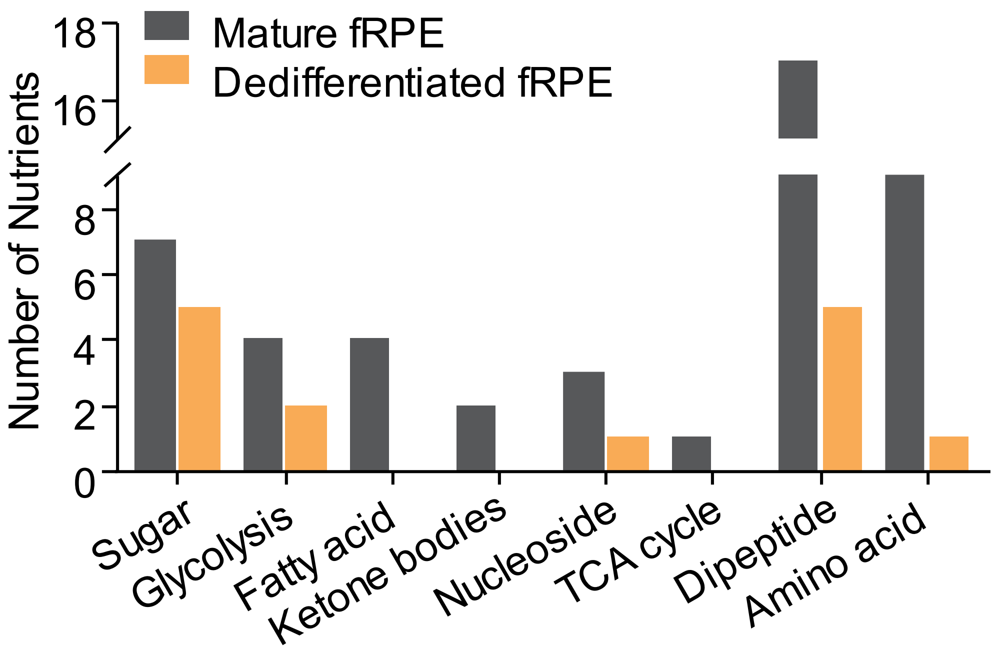


**Supplementary Figure S4. A comparison of metabolic phenotyping between mature fRPE and dedifferentiated fRPE**.

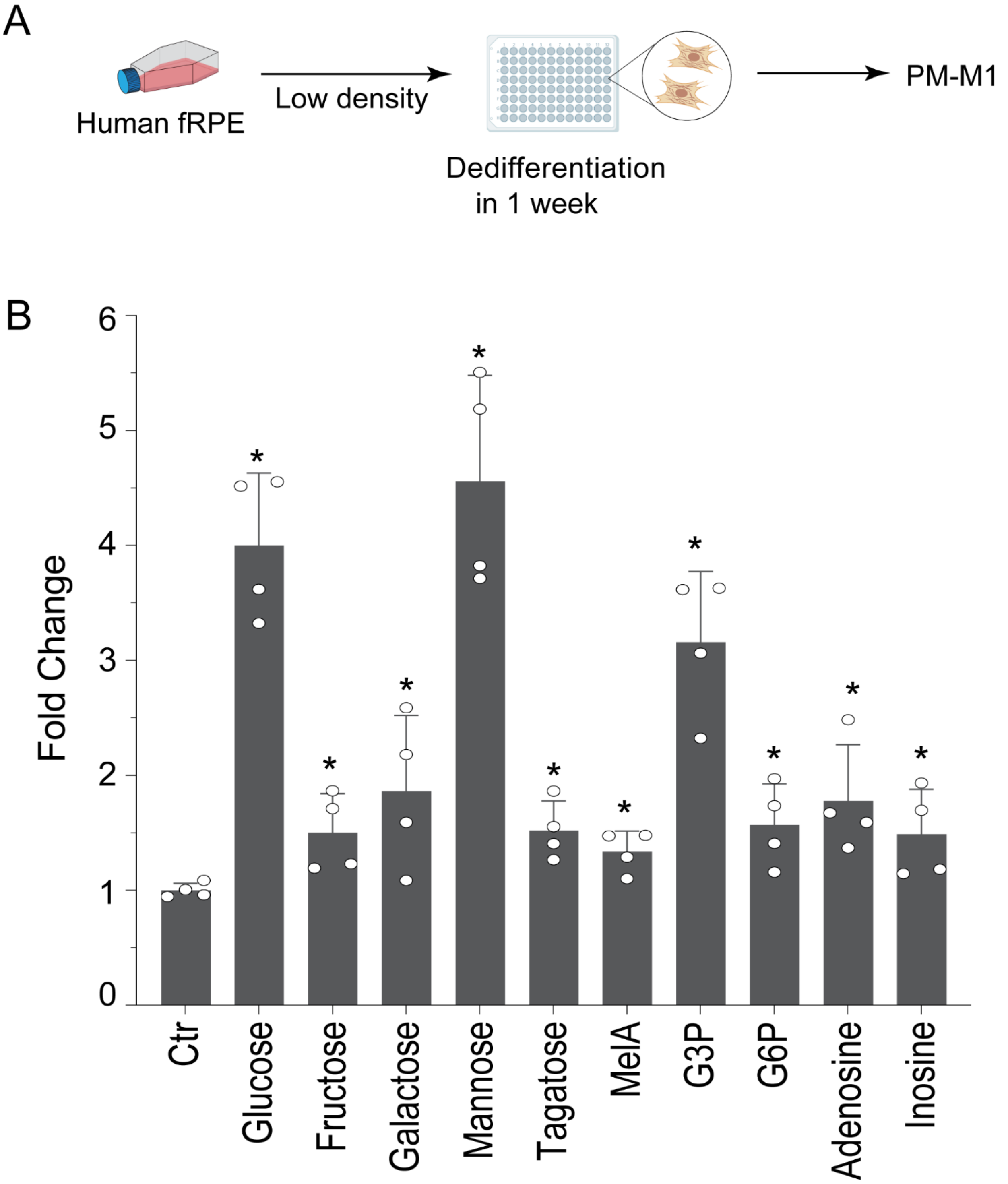


**Supplementary Figure S5. Nutrient utilization of dedifferentiated human fRPE cultured for 1 week.** (A) Human fRPE were seeded at 0.5 x $10^5$ cells per well and cultured for one week before metabolic phenotyping with PM-M1. (B) Nutrient utilization of dedifferentiated fRPE at 1 week. N =3. Fold change >1.5 or P<0.5 over Ctr.

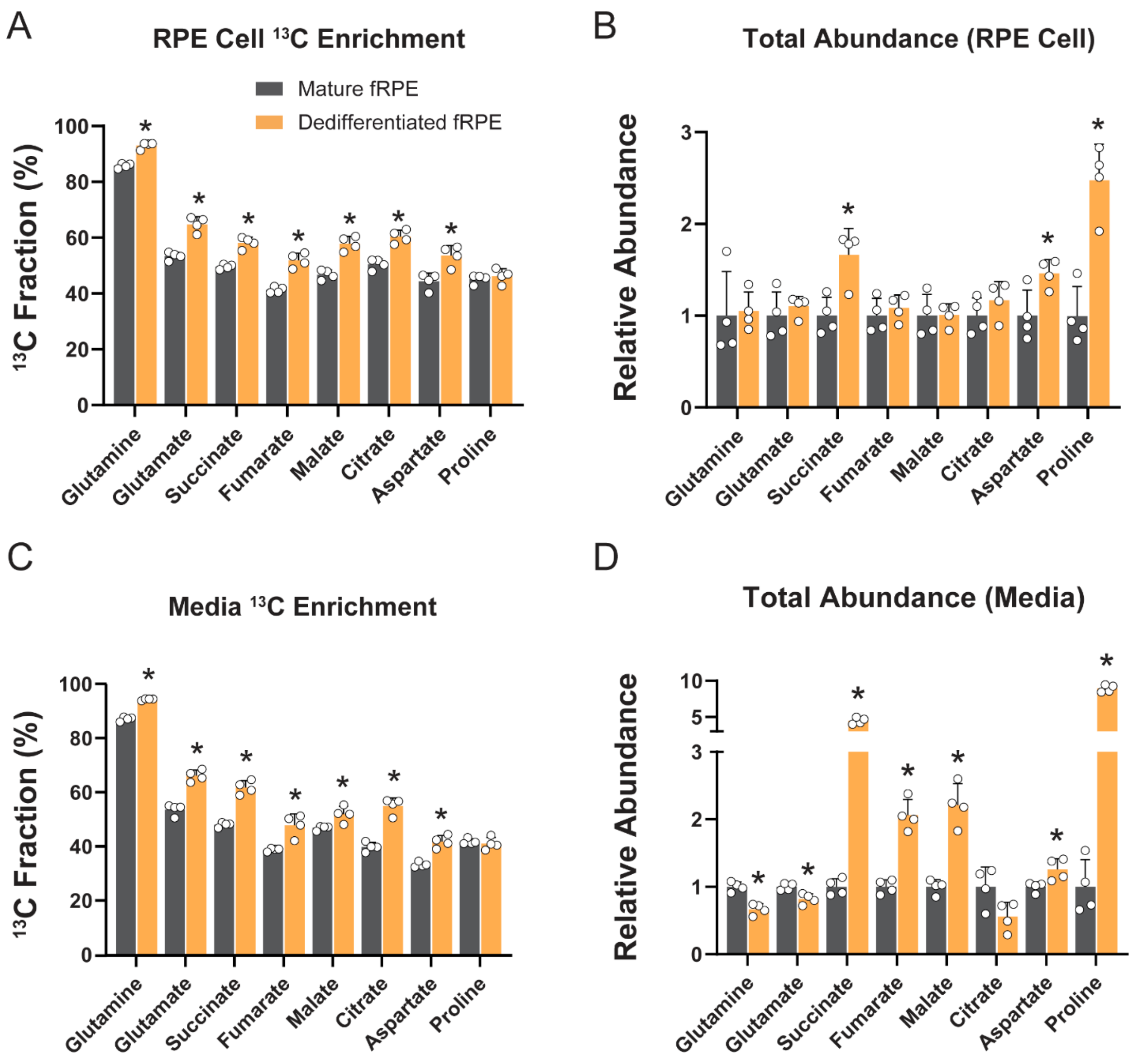


**Supplementary Figure S6. Dedifferentiated RPE cells enhance the utilization of $^{13}$C glutamine.** (A) The enrichment of $^{13}$C glutamine-derived metabolites in the media after incubation with 1 mM $^{13}$C glutamine for 48 hours in DMEM containing 5.5 mM glucose. (B) Total abundance of isotopologues in the culture media. Values were fold change relative to those in mature RPE. (C) The enrichment of $^{13}$C glutamine-derived metabolites in RPE cells. (D) Total abundance of isotopologues in RPE cells. N =4. *P<0.05 vs mature RPE.

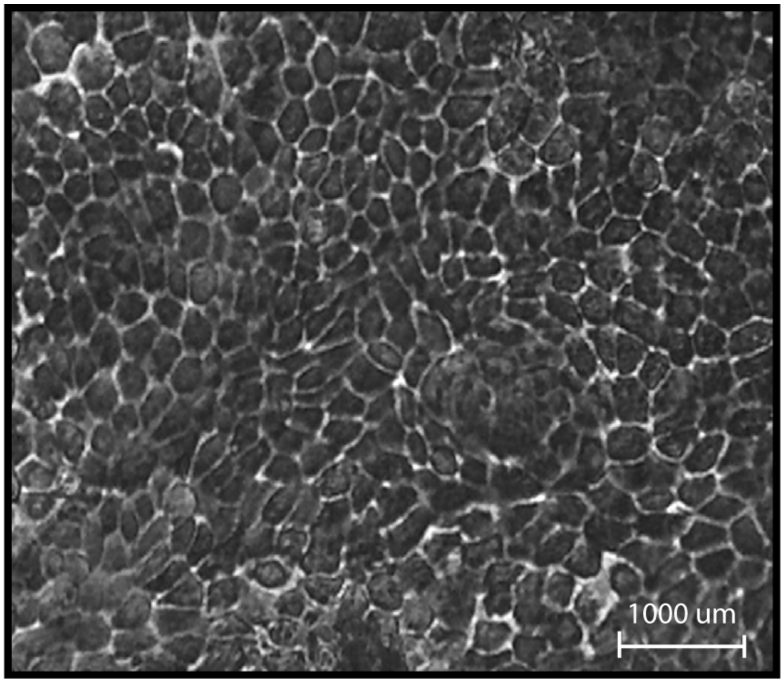


**Supplementary Figure S7. Morphology of human iPSC RPE cells**. A representative image of human iPSC RPE cells cultured for four weeks in 96-well plate before metabolic phenotyping. Scale = 1000µm.

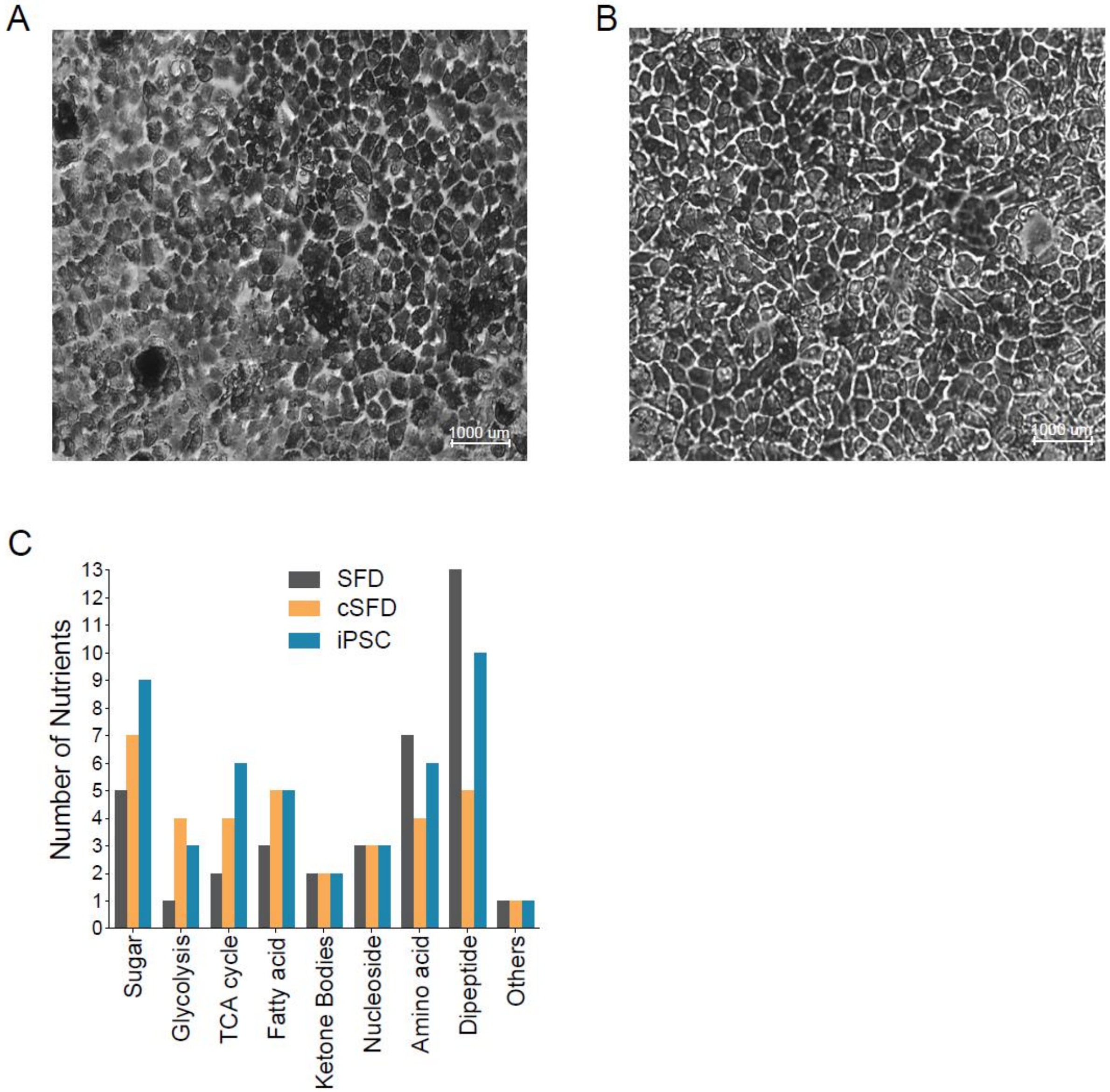


**Supplementary Figure S8. Metabolic phenotyping of Human iPSC SFD RPE cells and cSFD RPE cells**. (A-B). Representative bright-field images of human iPSC SFD RPE cells and cSFD RPE cells at 4 weeks before metabolic phenotyping. Scale = 1000µm. (C) A comparison of utilized nutrients by SFD RPE, cSFD RPE, and iPSC RPE

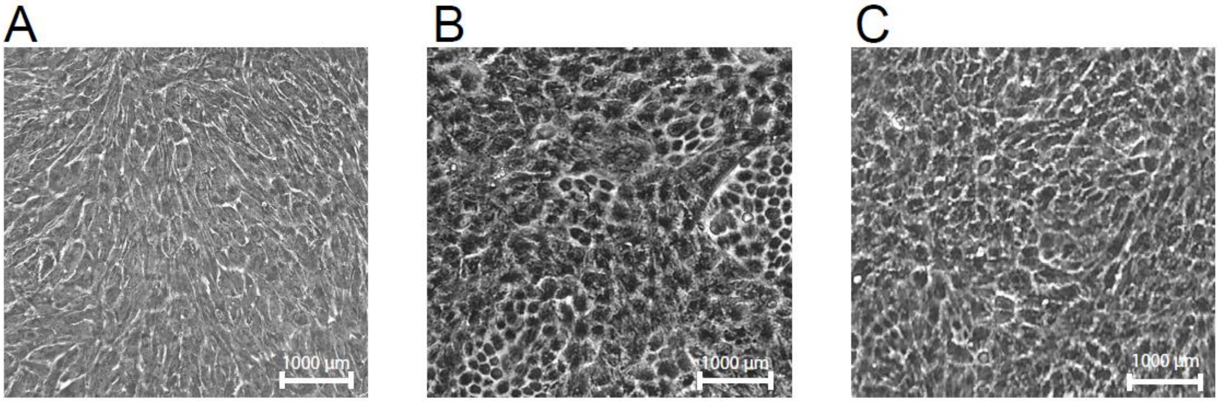


**Supplementary Figures S9. Metabolic phenotyping of human ARPE-19 cells cultured in different media**. Representative images of ARPE-19 cells cultured for 4 weeks in (A) DMEM/F-12 media, (B) MEM alpha RPE media, and (C) MEM alpha RPE media with 10 mM NAM. Scale = 1000µm

**Table S1 – Key Reagents and Materials.** All experimental materials used for this study are listed in this table.

| Reagent and Materials | Source | Identifier |
|---|---|---|
| **Biolog Screenings** | | |
| PM-M1 | Biolog | 13101 |
| PM-M2 | Biolog | 13102 |
| IF-M1 | Biolog | 72301 |
| Redox Dye Mix MA | Biolog | 74351 |
| FilterMax F5 Microplate Reader | Molecular Devices | 26 270-2064 |
| Softmax Pro 7.1 Software | Molecular Devices | 5049092 E |
| **Cell Culture** | | |
| DMEM/F12 | Thermo Fisher | 11320-033 |
| MEM Alpha | Thermo Fisher | 12561-072 |
| MEM Non-Essential Amino Acids Solution (100X) | Thermo Fisher | 11140-050 |
| Penicillin/Streptomycin | VWR | K952 |
| Fetal Bovine Serum (FBS) | VWR | 97068-085 |
| Dialyzed FBS | Cytiva Life Sciences | SH30079-03 |
| N1 Medium Supplement | Sigma | N6530 |
| Nicotinamide | Sigma | N3376 |
| Glutamine | Thermo Fisher | 25030081 |
| Dulbecco's Phosphate Buffered Saline 1X (DPBS) | VWR | 0119-0500 |
| Taurine | Sigma | T0625 |
| Hydrocortisone | Sigma | H6909 |
| 3,3',5-Triiodo-L-Thyronine | Sigma | T-5516 |
| Matrigel (Stem Cell Matrix) | Corning | 354234 |
| Trypsin-EDTA 0.25% (1X) | Thermo Fisher | 25200056 |
| Y-27632 Dihydrchloride (ROCK inhibitor) | SelleckChem | S1049 |
| CytoTune iPS 2.0 Sendai Reprogramming Kit | Thermo Fisher | A16517 |
| mTeSR 1 complete kit | Stemcell Technologies | 85850 |
| Amaxa Nucleofector Human Stem Cell Kit 2 | Thermo Fisher | VPH-5022 |
| 96-well Plate Sealing Tape | Thermo Fisher | 15036 |
| NalgeneTM Rapid-FlowTM Sterile Single Use Vacuum Filter Units | Thermo Fisher | 566-0020 |
| Tissue Culture Flask | Greiner Bio-One | 658170 |
| Reagent Reservoir | Thermo Fisher | 8096-11 |
| Tissue Culture Plate 96 wells | Thermo fisher | FB012931 |
| **Staining** | | |
| Phalloidin-568 | Invitrogen | A12380 |
| F-actin | Invitrogen | A12380 |
| αSMA | Abcam | ab7817 |
| Vimentin | Cell Signaling Tech | 5741 |
| Anti-Mouse Alexa Fluor 488 | Thermo Fisher | A-21202 |
| Anti-Rabbit Alexa Fluor 647 | Thermo Fisher | A-31573 |
| Fluoromount-G | SouthernBiotech | 0100-01 |

**Table S2 –** List of carbon nutrients in the PM-M1 microarray plates.

| **Well** | **Carbon Source** | **Abbr** | **Well** | **PM1 Carbon** | **Abbr** |
|---|---|---|---|---|---|
| A1-3 | Negative Control | Ctr | E3 | D-Galactose | Galactose |
| A4 | α-Cyclodextrin | | E4 | α-Methyl-D-Galactoside | |
| A5 | Dextrin | | E5 | β-Methyl-D-Galactoside | |
| A6 | Glycogen | | E6 | N-Acetyl-Neuraminic Acid | |
| A7 | Maltitol | | E7 | Pectin | |
| A8 | Maltotriose | | E8 | Sedoheptulosan | |
| A9 | D-Maltose | Maltose | E9 | Thymidine | |
| A10 | D-Trehalose | | E10 | Uridine | |
| A11 | D-Cellobiose | | E11 | Adenosine | |
| A12 | β-Gentiobiose | | E12 | Inosine | |
| B1 | D-Glucose-6-Phosphate | G6P | F1 | Adonitol | |
| B2 | α-D-Glucose-1-Phosphate | | F2 | L-Arabinose | |
| B3 | L-Glucose | | F3 | D-Arabinose | |
| B4-6 | α-D-Glucose | Glucose | F4 | β-Methyl-D-Xylopyranoside | |
| B7 | 3-O-Methyl-D-Mannoside | | F5 | Xylitol | |
| B8 | α-Methyl-D-Glucoside | | F6 | Myo-Inositol | Inositol |
| B9 | β-Methyl-D-Glucoside | | F7 | Meso-Erythritol | Meso-Ery |
| B10 | D-Salicin | | F8 | Propylene glycol | |
| B11 | D-Sorbitol | | F9 | Ethanolamine | |
| B12 | N-Acetyl-D-Glucosamine | GlcNAc | F10 | D, L-α-Glycerol Phosphate | G3P |
| C1 | D-Glucosaminic Acid | | F11 | Glycerol | |
| C2 | D-Glucuronic Acid | | F12 | Citric Acid | Citrate |
| C3 | Chondroitin-6-Sulfate | | G1 | Tricarballylic Acid | Tcba |
| C4 | Mannan | | G2 | D, L-Lactic Acid | Lactate |
| C5 | D-Mannose | Mannose | G3 | Methyl D-lactate | |
| C6 | α-Methyl-D-Mannoside | | G4 | Methyl pyruvate | mPyr |
| C7 | D-Mannitol | | G5 | Pyruvic Acid | Pyruvate |
| C8 | N-Acetyl-β-D-Mannosamine | | G6 | α-Ketoglutaric acid | α-KG |
| C9 | D-Melezitose | | G7 | Succinamic Acid | SucA |
| C10 | Sucrose | | G8 | Succinic Acid | Succinate |
| C11 | Palatinose | | G9 | Mono-Methyl Succinate | mmSuc |
| C12 | D-Turanose | Turanose | G10 | L-Malic Acid | L-Mal |
| D1 | D-Tagatose | Tagatose | G11 | D-Malic Acid | |
| D2 | L-Sorbose | | G12 | Meso-Tartaric Acid | Meso-Tar |
| D3 | L-Rhamnose | | H1 | Acetoacetic Acid (alpha) | AcAc |
| D4 | L-Fucose | | H2 | γ-Amino-N-Butyric Acid | GABA |
| D5 | D-Fucose | | H3 | α-Ketobutyric acid | α-KBA |
| D6 | D-Fructose-6-Phosphate | | H4 | α-Hydroxy Butyric acid | α-HBA |
| D7 | D-Fructose | Fructose | H5 | D, L- β-Hydroxy Butyric Acid | 3HB |
| D8 | Stachyose | | H6 | γ-Hydroxy Butyric Acid | γ-HBA |
| D9 | D-Raffinose | | H7 | Butyric Acid | Butyrate |
| D10 | D-Lactitol | | H8 | 2,3-Butanediol | |
| D11 | Lactulose | | H9 | 3-Hydroxy-2-Butanone | |
| D12 | α-D-Lactose | Lactose | H10 | Propionic Acid | Propionate |
| E1 | Melibionic Acid | MelA | H11 | Acetic Acid | Acetate |
| E2 | D-Melibiose | | H12 | Hexanoic Acid | Hexanoate |

* Abbr, Abbreviation

**Table S3** - List of nitrogen nutrients in PM-M2 microarray plates.

| Well | Nitrogen Source | Abbr | Well | Nitrogen Source | Abbr |
|---|---|---|---|---|---|
| A1-3 | Negative Control | Ctr | E2 | Alanine-Serine | Ala-Ser |
| A4 | Tween 20 | T20 | E3 | Alanine-Threonine | Ala-Thr |
| A5 | Tween 40 | T40 | E4 | Alanine-Tryptophan | Ala-Trp |
| A6 | Tween 80 | T80 | E5 | Alanine-Tyrosine | Ala-Tyr |
| A7 | Gelatin | | E6 | Alanine-Valine | Ala-Val |
| A8 | L-Alaninamide | AlaN | E7 | Arginine-Alanine | Arg-Ala |
| A9 | L-Alanine | Ala | E8 | Arginine-Arginine | Arg-Arg |
| A10 | D-Alanine | D-Ala | E9 | Arginine-Asparagine | Arg-Asp |
| A11 | L-Arginine | Arg | E10 | Arginine-Glutamine | Arg-Gln |
| A12 | L-Asparagine | Asn | E11 | Arginine-Glutamate | Arg-Glu |
| B1 | L-Aspartic Acid | Asp | E12 | Arginine-Isoleucine | Arg-Ile |
| B2 | D-Aspartic Acid | D-Asp | F1 | Arginine-Leucine | Arg-Leu |
| B3 | L-Glutamic Acid | Glu | F2 | Arginine-Lysine | Arg-Lys |
| B4 | D-Glutamic Acid | D-Glu | F3 | Arginine-Methionine | Arg-Met |
| B5 | L-Glutamine | Gln | F4 | Arginine-Phenylalanine | Arg-Phe |
| B6 | Glycine | Gly | F5 | Arginine-Serine | Arg-Ser |
| B7 | L-Histidine | His | F6 | Arginine-Tryptophan | Arg-Trp |
| B8 | L-Homoserine | | F7 | Arginine-Tyrosine | Arg-Tyr |
| B9 | Hydroxy-L-Proline | | F8 | Arginine-Valine | Arg-Val |
| B10 | L-Isoleucine | Ile | F9 | Asparagine-Glutamate | Asn-Glu |
| B11 | L-Leucine | Leu | F10 | Asparagine-Valine | Asn-Val |
| B12 | L-Lysine | Lys | F11 | Aspartate-Alanine | Asp-Ala |
| C1 | L-Methionine | Met | F12 | Aspartate-Aspartate | Asp-Asp |
| C2 | L-Ornithine | Orn | G1 | Aspartate-Glutamate | Asp-Glu |
| C3 | L-Phenlyalanine | Phe | G2 | Aspartate-Glutamine | Asp-Gln |
| C4 | L-Proline | Pro | G3 | Aspartate-Glycine | Asp-Gly |
| C5 | L-Serine | Ser | G4 | Aspartate-Leucine | Asp-Leu |
| C6 | D-Serine | D-Ser | G5 | Aspartate-Lysine | Asp-Lys |
| C7 | L-Threonine | Thr | G6 | Aspartate-Phenylalanine | Asp-Phe |
| C8 | D-Threonine | D-Thr | G7 | Aspartate-Tryptophan | Asp-Trp |
| C9 | L-Tryptophan | Trp | G8 | Aspartate-Valine | Asp-Val |
| C10 | L-Tyrosine | Tyr | G9 | Glutamate-Alanine | Glu-Ala |
| C11 | L-Valine | Val | G10 | Glutamate-Aspartate | Glu-Asp |
| C12 | Alanine-Alanine | Ala-Ala | G11 | Glutamate-Glutamate | Glu-Glu |
| D1 | Alanine-Arginine | Ala-Arg | G12 | Glutamate-Glycine | Glu-Gly |
| D2 | Alanine-Asparagine | Ala-Asn | H1 | Glutamate-Serine | Glu-Ser |
| D3 | Alanine-Aspartate | Ala-Asp | H2 | Glutamate-Tryptophan | Glu-Trp |
| D4 | Alanine-Glutamate | Ala-Glu | H3 | Glutamate-Tyrosine | Glu-Tyr |
| D5 | Alanine-Glutamine | Ala-Gln | H4 | Glutamate-Valine | Glu-Val |
| D6 | Alanine-Glycine | Ala-Gly | H5 | Glutamine-Glutamate | Gln-Glu |
| D7 | Alanine-Histidine | Ala-His | H6 | Glutamine-Glutamine | Gln-Gln |
| D8 | Alanine-Isoleucine | Ala-Ile | H7 | Glutamine-Glycine | Gln-Gly |
| D9 | Alanine-Leucine | Ala-Leu | H8 | Glycine-Alanine | Gly-Ala |
| D10 | Alanine-Lysine | Ala-Lys | H9 | Glycine-Arginine | Gly-Arg |
| D11 | Alanine-Methionine | Ala-Met | H10 | Glycine-Asparagine | Gly-Asn |
| D12 | Alanine-Phenylalanine | Ala-Phe | H11 | Glycine-Aspartate | Gly-Asp |
| E1 | Alanine-Proline | Ala-Pro | H12 | α-D-Glucose | D-Glucose |

* Abbr, Abbreviation
